# Alterations in platelet proteome signature and impaired platelet integrin α_IIb_β_3_ activation in patients with COVID-19

**DOI:** 10.1101/2022.10.12.511145

**Authors:** Lucy J. Goudswaard, Christopher M. Williams, Jawad Khalil, Kate L. Burley, Fergus Hamilton, David Arnold, Alice Milne, Phil A. Lewis, Kate J. Heesom, Stuart J. Mundell, Andrew D. Davidson, Alastair W. Poole, Ingeborg Hers

## Abstract

**Background:** Patients with coronavirus disease-19 (COVID-19) are at increased risk of thrombosis, which is associated with altered platelet function and coagulopathy, contributing to excess mortality.

**Objectives:** We aimed to characterise the mechanism of altered platelet function in COVID-19 patients.

**Methods:** The platelet proteome, platelet functional responses and platelet-neutrophil aggregates were compared between patients hospitalised with COVID-19 and healthy control subjects using Tandem Mass Tag (TMT) proteomic analysis, Western blotting and flow cytometry.

**Results:** COVID-19 patients showed a different profile of platelet protein expression (858 altered out of 5773 quantified). Levels of COVID-19 plasma markers were enhanced in COVID-19 platelets. Gene ontology (GO) pathway analysis demonstrated that levels of granule secretory proteins were raised, whereas some platelet activation proteins, such as the thrombopoietin receptor and PKCα, were lowered. Basally, COVID-19 platelets showed enhanced phosphatidylserine (PS) exposure, with unaltered integrin α_IIb_β_3_ activation and P-selectin expression. Agonist-stimulated integrin α_IIb_β_3_ activation and PS exposure, but not P-selectin expression, were significantly decreased in COVID-19 patients. COVID-19 patients had high levels of platelet-neutrophil aggregates, even under basal conditions, compared to controls. This interaction was disrupted by blocking P-selectin, demonstrating that platelet P-selectin is critical for the interaction.

**Conclusions:** Overall, our data suggests the presence of two platelet populations in patients with COVID-19: one with circulating platelets with an altered proteome and reduced functional responses and another with P-selectin expressing neutrophil-associated platelets. Platelet driven thromboinflammation may therefore be one of the key factors enhancing the risk of thrombosis in COVID-19 patients.

**Essentials:**

- COVID-19 patient platelet function and platelet proteins were compared with healthy controls
- Proteomic analysis of platelets indicated that COVID-19 decreased platelet activation proteins
- Agonist induced PS exposure and integrin α_IIb_β_3_ activation were impaired in COVID-19
- COVID-19 led to maximal levels of P-selectin dependent platelet-neutrophil aggregates

## Introduction

The Coronavirus disease-19 (COVID-19) pandemic started in China in 2019 and rapidly spread over the world causing over 6.5 million deaths to date [1]. The disease is caused by a single positive sense RNA virus; severe acute respiratory syndrome coronavirus 2 (SARS-CoV-2) and can cause a wide variety of clinical syndromes, with most patients hospitalised with pneumonitis. During the COVID-19 pandemic it has become apparent that patients with COVID-19 have enhanced rates of venous and arterial thrombosis, including deep-vein thrombosis, pulmonary embolism, myocardial infarction, and ischemic stroke [2–6]. Circulating platelet microaggregates were detected in COVID-19 patients, showing a strong correlation to COVID-19 severity [7, 8]. Post-mortem examinations revealed microthrombi in the lung, kidney, liver, heart and brain, suggesting COVID-19 can induce systemic thrombosis, thereby contributing to multi-organ failure [9–14]. COVID-19 is also associated with elevated serum coagulation markers such as fibrinogen and D-dimer, with the latter related to disease severity and poor prognosis [2, 15, 16]. Thromboprophylaxis, such as administration of heparin, has been shown to improve outcomes in hospitalised COVID-19 patients, but had a limited effect on mortality rates of the subset of patients with severe COVID-19 [17–20]. Thrombosis thus remains a prominent feature of COVID-19 despite thromboprophylaxis, suggesting that the picture is complex, with raised thrombin generation not the only contributing factor.

Major players in thrombosis are platelets, blood cells essential for haemostasis but that also contribute to thrombosis when inappropriately activated. Data obtained during the COVID-19 pandemic suggest that platelets become hyperactive, with reports of increased secretion of dense and α-granules, increased aggregation and increased formation of platelet-leukocyte aggregates [21–23]. Platelet transcriptome analysis showed an overrepresentation of pathways involved in antigen presentation and mitochondrial dysfunction in COVID-19 patients, potentially contributing to platelet hyperactivity [21]. However, there are also reports of impaired or reduced platelet functional responses in COVID-19 patients [24, 25], suggesting that the platelet response is complex.

The pathogenesis of thrombosis in COVID-19 patients is not completely understood, but it has many hallmarks of thromboinflammation [2, 26]. It is likely that SARS-CoV-2 activates endothelial cells via the angiotensin converting enzyme 2 (ACE2) receptor [27], thereby causing vascular dysfunction [28]. Damaged endothelium leads to activation of the innate immune system mediated through complement, pro-inflammatory cytokines, tissue factor (TF) expression and neutrophil recruitment [2, 28–31].

Together, this results in upregulation of adhesion molecules and enhances the expression and/or release of prothrombotic factors such as neutrophil extracellular traps (NETS) and von Willebrand factor (vWF). The latter will bind and activate platelets, leading not only to aggregate and thrombus formation but also feed back to further activation of the innate immune system by the release of cytokines/chemokines and direct platelet-leukocyte interactions [14]. Platelets then amplify thrombin generation through the expression of tissue factor (TF) and phosphatidyl serine [2, 32], further promoting platelet activation and the cleavage of fibrinogen into the fibrin network required for clot formation.

This intricate network of interactions between platelets, the innate immune system and the coagulation cascade are likely factors contributing to thrombosis in COVID-19 patients [26]. As well as indirect activation via damaged endothelium, it is also hypothesised that SARS-CoV-2 directly interacts with platelets, through receptors such as TLR7, FcγRIIA and CD147, evidenced to contribute to platelet hyperactivity and thrombosis as well [33–35]. Platelet expression of ACE2, a receptor involved in SARS-CoV-2 binding and cellular infection, is controversial; although a few studies reported platelet ACE2 expression [36], the majority failed to detect ACE2 by western blotting, transcriptomics and proteomics [37–39].

Although the emphasis has been predominantly on coagulopathy and activation of the immune system as important precipitants of thrombosis in patients with COVID-19, it more recently has become apparent that platelets are important, but poorly understood, drivers in this process [40]. In this study, we therefore aimed to assess the effect of COVID-19 on the platelet proteome and relate this to platelet functional responses and platelet-neutrophil aggregate formation in patients hospitalised with COVID-19.

## Methods

### Study population

Twenty adult patients hospitalised with COVID-19 from the DISCOVER study at Southmead Hospital, Bristol, UK, and nineteen healthy control adult participants (from the University of Bristol, UK) were enrolled in the study between October 2020 and February 2021 after providing written informed consent. A detailed description of the DISCOVER cohort is available elsewhere [41] but briefly consecutive patients presenting to hospital with suspected or RT-PCR confirmed COVID-19 were recruited. Demographics, comorbidities, medications, treatment records, and admission results were recorded. Control participants did not have COVID-19 (self-reported) and were matched for blood draw time. Control participants were not recruited if they were pregnant, taking antiplatelet drugs or had known clotting or bleeding disorders, or a blood-borne disease. See the supplementary methods section for further details. Full blood counts were measured in COVID-19 patients and healthy controls using the Sysmex XN-20 haematology analyser.

### Ethics

Approval of this study was granted by the Yorkshire & The Humber - South Yorkshire Research Ethics Committee (NHS-REC reference 20/YH/0121) and South Central - Hampshire A Research Ethics Committee (NHS-REC reference 20/SC/0222).

### Isolation of platelet rich plasma (PRP) and washed platelets

Venous blood was taken by venepuncture into 4.5 mL vacutainers containing 3.2% sodium citrate (1:7 v/v) and centrifuged (1000 RPM, 17 mins) to produce PRP. For washed platelets, PRP was acidified with 1:7 v/v acidic citrate dextrose before a second centrifugation (1700 RPM, 10 mins). Platelets were double washed in modified CGS buffer (120 mM NaCl, 25.8 mM sodium citrate dihydrate, 0.1% [w/v] D-glucose, 0.02 U mL^−1^, pH 6.5) followed by centrifugation (1700 RPM, 10 mins). Washed platelets were resuspended in modified HEPES-Tyrode’s buffer (145 mM NaCl, 3 mM KCl, 0.5 mM Na_2_HPO_4_, 1 mM MgS0_4_.7H_2_O, 10 mM HEPES, pH 7.4, 0.1% [w/v] D-glucose) with 0.02 U mL^−1^ apyrase).

### Preparation of platelet samples for tandem mass tag (TMT) proteomic analysis

Washed platelets were lysed with radioimmunoprecipitation assay buffer (RIPA: 25 mM HEPES, 200 mM NaCL, 1mM EDTA, 1 % (v/v) NP40, 0.5 % (w/v) sodium deoxychelate, 0.1 % (w/v) SDS) supplemented with cOmplete™ Mini and PhosSTOP™. Samples were cleared by centrifugation at 4 °C, heated to 55 °C and stored at −80 °C. 50 μg of sample was digested with trypsin, labelled with TMT eleven plex reagents and subjected to TMT analysis as described in the Supplementary methods section.

### Comparison of platelet proteome in COVID-19 and healthy control subjects

A principal component analysis (PCA) was used to identify and remove outliers. Normalised protein abundances were log_2_ transformed. The fold change was calculated by subtracting the mean abundance of the control subjects from the mean protein abundance of the COVID-19 patients. Student’s unpaired two-tailed t-tests were used to determine differences in protein levels. Multiple testing was accounted for using a false discovery rate (FDR) p-value correction. A positive log_2_ fold change can be interpreted as an increase in protein levels in COVID-19 patients. Strength of association was determined by a composite of the log_2_ fold change in protein levels and the p-value.

### Exploring platelet pathway changes using gene ontology (GO) pathways

We explored whether plasma or serum proteins reported to be affected by COVID-19 were also affected in platelets [42–44]. To explore the effect that a COVID-mediated change in platelet proteins may have on platelet function, we integrated proteomic results with GO pathway annotations. This included proteins involved in platelet secretion (alpha- and dense-granule lumen gene products i.e. GO pathways GO:0031093 and GO:0031089 respectively) and platelet activation. The latter was explored using seven GO pathways: negative regulation of platelet activation (GO:0010544), negative regulation of platelet aggregation (GO:0090331), negative regulation of platelet aggregation in another organism (GO:0035893), positive regulation of platelet activation (GO:0010572), positive regulation of platelet aggregation (GO:1901731), regulation of platelet activation (GO:0010543) and regulation of platelet aggregation (GO:0090330).

### Western blotting

Washed platelets were lysed in NuPage LDS sample buffer supplemented with 1:10 v/v dithiothreitol (DTT) followed by Tris-Glycine sodium dodecyl sulfate-polyacrylamide gel electrophoresis (10% SDS-PAGE) and immunoblotted for the TPO receptor (c-Mpl), PKCa, RalB, PKA-RIIb, IFITM3 and GAPDH as described previously [45].

### Flow cytometry analysis

Washed platelets, PRP and whole blood were diluted in HEPES-Tyrode’s buffer (2×10^7^ washed platelets/mL, PRP (1:40 v/v) and whole blood (1:10 v/v)). Platelets were incubated with agonists (concentration-response curves for PAR1-AP, ADP or collagen-related peptide (CRP)) or costimulated with 1 U/mL thrombin and 10 ug/mL CRP for 10 min where indicated. To measure integrin α_IIb_β_3_ activation and P-selectin expression, PRP or whole blood was incubated with FITC-conjugated PAC1 or PE-conjugated CD62P antibodies respectively (1:10 (v/v)). Platelet receptor levels were explored using FITC or PE-conjugated antibodies in PRP. PS exposure was measured using Annexin V in washed platelets. Samples were fixed with 1x FixLyse (3:1 v/v, whole blood only), followed by a final concentration of 4% PFA (PRP, whole blood), or immediately quenched with HEPES-Tyrode’s buffer containing 2 mM CaCl_2_ (4:1 v/v, washed platelet PS exposure only). A total of 10,000 platelet events were collected on a BD Accuri™ C6 Plus flow cytometer (BD Biosciences, Wokingham, UK) in the platelet FSC/SSC gate (washed platelets, PRP) and FSC/SSC gate/CD42b (whole blood). The neutrophil population in whole blood was gated on CD45 and SSC parameters with 1000 neutrophil events captured. Data is expressed as the Median Fluorescence Intensity (FMI, PAC1 and P-selectin), % of positive platelets (annexin V) or as % of neutrophils associated with platelets (CD41+/CD45+events).

### Platelet function data

If data met the assumption of normality, as tested by a Shapiro-Wilk test, data was analysed using parametric tests (unpaired, two-tailed student t-test), otherwise a nonparametric test was used (Mann-Whitney test). Concentration-response curves were fitted using a four-parameter logistic curve fit. Differences in curve fit were explored by comparing the individual logEC50s and curve maximum values of each group. P-values are reported throughout where P<0.05, as a guidance for sufficient evidence to reject the null hypothesis. Analyses were performed using Prism 8 or R version 4.0.2 [46].

### See supplementary information for Materials and TMT details

## Results

### Participant characteristics

The mean age of the recruited COVID-19 cohort was 59 years (SD of 15.7 years), whereas the healthy control donor cohort recruited were younger, with a mean age of 39 years (SD 13.8) (p=0.0002, Table 1). Control and COVID-19 groups included a similar proportion of males and females. The mean BMI for COVID-19 patients was 30.6 kg/m^2^ (SD 4.4 kg/m^2^) compared to 22.7 (SD 2.8kg/m^2^) for controls (p<0.0001). Patients were supported with oxygen therapy (88 %) with additional treatments including heparin (95 %), dexamethasone (89 %) and rivaroxaban (6 %) (Table 1).

**Table 1.** Characteristics of study participants.

| Variable | | Control group mean (±SD) or % | Control group N | COVID-19 group mean (±SD) or % | COVID-19 group N | P value for difference |
| --- | --- | --- | --- | --- | --- | --- |
| **Age (years)** |  | 39.0 (13.8) | 18 | 59.1 (15.7) | 19 | 0.0002 |
| **Sex** | Male | 63 % | 19 | 68 % | 19 | 0.73* |
|  | Female | 37 % |  | 32 % |  |  |
| **BMI (kg/m<sup>2</sup>)** |  | 22.7 (2.8) | 18 | 30.6 (4.4) | 15 | <0.0001 |
| **Oxygen required (N)** |  | NA | NA | 88 % | 16 | NA |
| **Medication** | Heparin | NA | NA | 95 % | 18 | NA |
|  | Dexamethasone |  |  | 89 % |  |  |
|  | Rivaroxaban |  |  | 6 % |  |  |

\*=Chi squared test

### Full blood count participants

COVID-19 patients had elevated neutrophil counts (6.6 × 10^9^/L ± SD 3.5 vs 2.6 × 10^9^/L ± SD 0.8, p=6×10^−5^), but a reduced lymphocyte count (1.1 × 10^9^/L ± SD 1 vs 1.8 × 10^9^/L ± SD 0.4, p=0.01). Platelet parameters (platelet count, immature platelet fraction and immature platelet count) were unchanged (S.Table 1).

### The platelet proteome signature is altered in COVID-19 patients

To study whether SARS-CoV-2 viral infection alters the platelet proteome, we performed TMT labeling on lysates of washed platelet samples from seven COVID-19 patients and six healthy controls. After accounting for multiple testing using an FDR p-value adjustment, 858 proteins out of 5773 were altered in abundance in platelet lysates from patients with COVID-19 compared to healthy controls (FDR p < 0.05, 14.9 % of total proteins detected, Figure 1a, STable.2). Proteins that increased in amount in platelets from COVID-19 patients compared to healthy controls include the 40S ribosomal proteins S5 and S3 with log_2_ fold changes of 1.07 (p = 0.001) and 1.11 (p = 0.003), respectively and C-reactive protein (log_2_ fold change 3.55, p = 0.003). We also found weak evidence for an increase in the amounts of the antiviral immune protein interferon-induced transmembrane membrane protein 3 (IFITM3) in platelets from COVID-19 patients compared to healthy controls, with a log_2_ fold change of 2.02 (p = 0.096), however this did not pass the FDR multiple testing threshold. Interestingly, multiple platelet activation signaling proteins were decreased in platelets from COVID-19 patients compared to healthy controls, including the thrombopoietin (TPO) receptor cMpl (log_2_ fold change of −1.45, p = 4.8×10^−4^) and protein Kinase C α (PKCα) (log_2_ fold change −0.60, p = 0.008).

**Figure 1.**
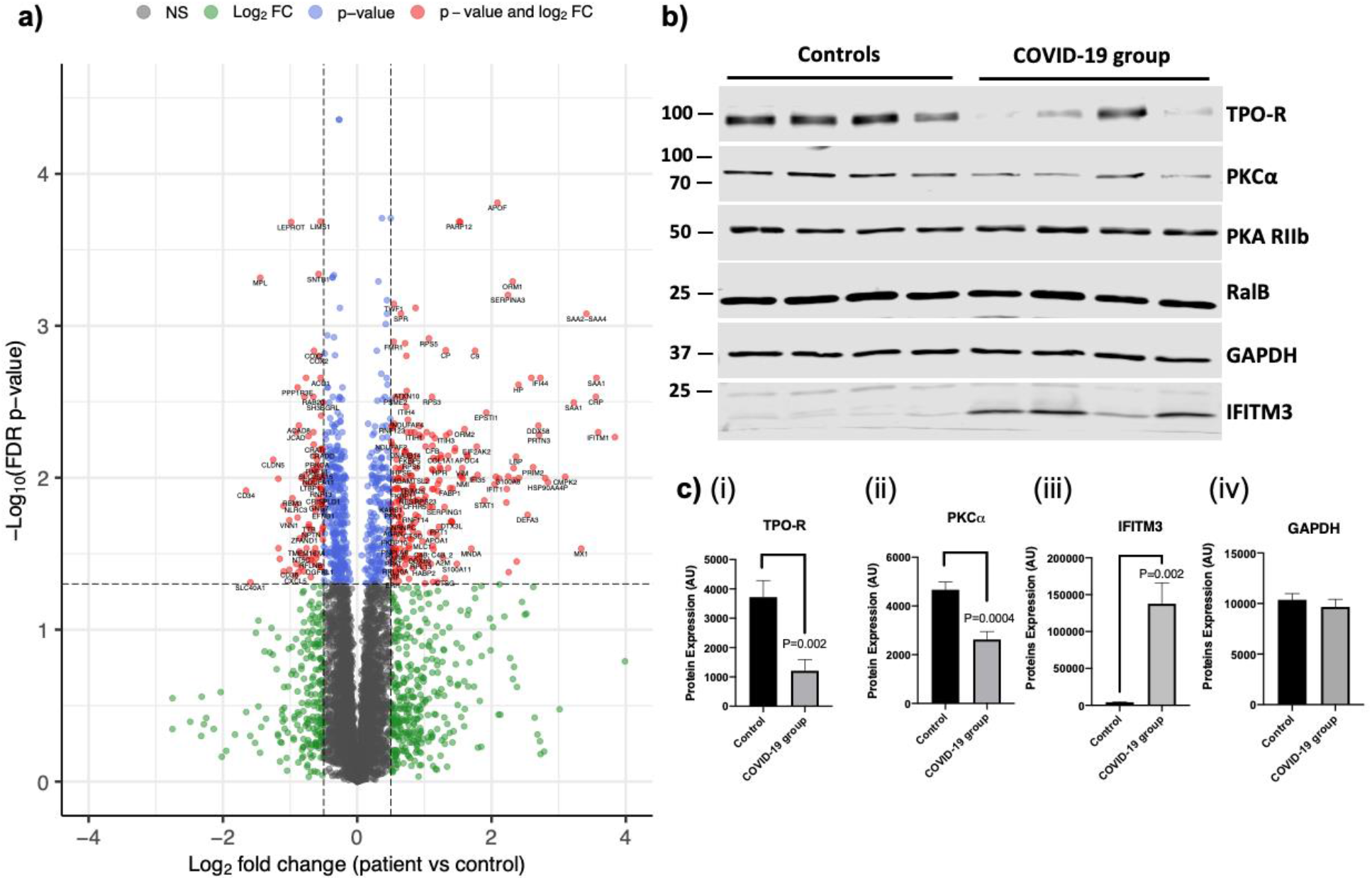
The platelet proteome is altered in patients with COVID-19. a) Volcano plot of the proteins altered in platelet lysates from patients (N=7) and controls (N=6) using TMT mass spectrometry. Protein changes were analysed by a Student’s t-test. The proteins that had both a larger log_2_(fold change) than ± 0.5 and a smaller false discovery rate (FDR) adjusted P-value than 0.05 are shown in red. Proteins which only had a larger log_2_(fold change) than ± 0.5 are shown in green. Proteins with only a smaller FDR P-value than 0.05 are shown in blue. Proteins that did not reach the P-value or log_2_(fold change) threshold are shown in grey. One protein was excluded from the figure (which had a logFC of over −4) to be able to see other data points more clearly. Plot made using the Enhanced Volcano R package https://github.com/kevinblighe/EnhancedVolcano). b) Representative Western blot using platelet lysates from four controls and four patients for the TPO receptor, PKCα, PKA (control), RalB, GAPDH (control) and IFITM3. Analyses performed a using Student’s t-test c) (i) Western blot quantification of the TPO receptor (mean + SEM, N=9). (ii) Western blot quantification of PKCα (mean + SEM, N=9). (iii) Western blot quantification of IFITM3 protein expression (mean + SEM, N=8). (iv) Western blot quantification of GAPDH loading control (mean + SEM, N=8, p=0.58).

To confirm the untargeted TMT proteomic results, we next performed Western blotting analysis on platelet lysates from at least eight COVID-19 patients and eight healthy controls. Platelet lysates from the COVID-19 group again showed a reduction in the expression of the TPO receptor cMpl (1211 ± 567.4 vs 3719 ± 382, p = 0.002) and PKCα (2629 ± 323.5 vs 4688 ± 319.2, p = 0.0004) compared to healthy controls (Figures 1b, 2ci, 2cii). In contrast, protein kinase A (PKA) expression levels were unaltered, in agreement with our proteomic findings. We also found evidence for an increase in levels of IFITM3 in platelet lysates from COVID-19 patients compared to healthy controls (Figure 1b, Figure 1ciii), as found in a previous study [21]. Interestingly, proteomic analysis failed to detect any of the SARS-CoV-2 viral proteins reported by Davidson et al. [47] in COVID-19 patient samples, indicating that the virus had not entered and/or replicated in platelets. The receptor for SARS-CoV2, ACE2, was also not detected in any sample. In contrast, CD147 (basigin), a receptor with suggested interaction with SARS-CoV2 [48], was present in both control and patient samples and was unaltered in the COVID-19 group (S.Table 2).

### Enhanced platelet content of COVID-19 plasma biomarkers

Previous studies have identified multiple plasma or serum proteins associated with COVID-19 [42–44]. Platelets are not only able to release their granule content but can also selectively take up proteins from their environment [49]. We were therefore interested in whether the platelet content of these plasma/serum biomarkers is altered in COVID-19 patients. Using TMT proteomic analysis, we detected 38 proteins that have been demonstrated to be associated with COVID-19 in plasma/serum in at least two out of three studies[42–44]. Of these, 12 proteins were altered in COVID-19 patients’ platelets (Figure 2a) compared to controls. Interestingly, except for gelsolin, levels of 11 platelet plasma biomarkers were enhanced in COVID-19 platelets, as shown in the waterfall plot in Figure 2a. Of these, four proteins showed more than a four-fold increase (log_2_ fold change of >2) compared to control platelets; serum amyloid A (SAA1), C-reactive protein (CRP), lipopolysaccharide binding protein (LBP) and galectin 3-binding protein (G3BP). These results suggest that platelets take up a subset of COVID-19 associated plasma proteins.

**Figure 2.**
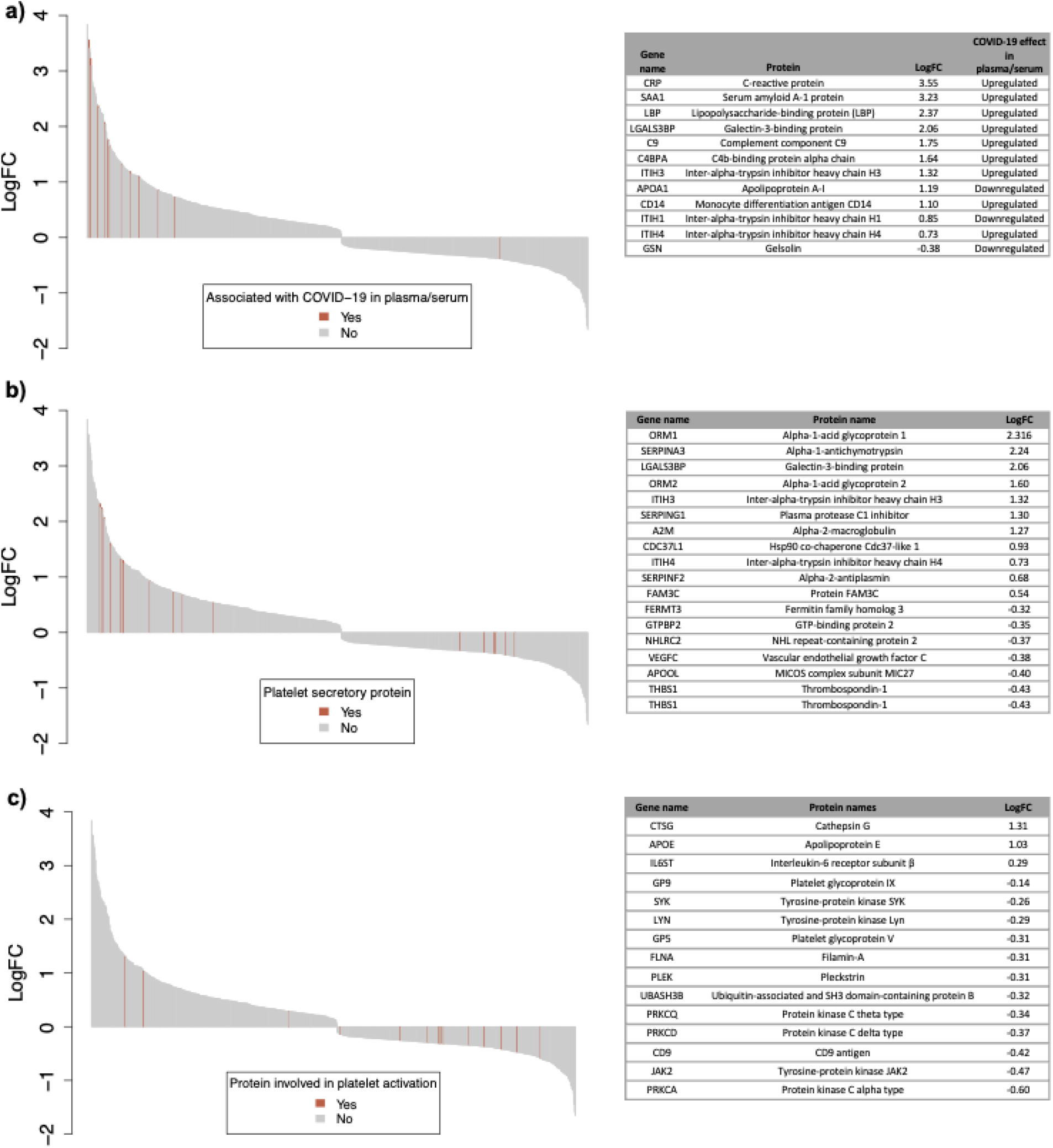
Waterfall plots displaying the log2 fold change of platelet proteins in COVID-19 patients compared to controls. Each protein is represented by a grey bar and proteins in specific pathways are indicated with red lines. a) Proteins highlighted in red and listed in the table have been shown to be associated with COVID-19 in plasma or serum in at least two studies by Messner et al. [42], Leng et al. [43] or Roh et al. [44] b) Proteins highlighted in red and listed in the table have known involvements in gene ontology (GO) pathways alpha- and dense-granule lumen gene products (GO:0031093 and GO:0031089 respectively) c) Proteins highlighted in red and listed in the table have known involvements in platelet activation or platelet aggregation GO pathways.

### Alterations in proteins involved in platelet secretion pathways in COVID-19 platelets

To investigate whether proteins involved in granule secretion were altered in platelets from COVID-19 patients compared to healthy controls, we compared changes in proteins using GO pathway annotations [50, 51]. A total of 70 (68 unique) proteins were involved in the α-granule and dense granule lumen gene product GO pathways and detected in the current study. Of these 18 (17 unique) proteins were associated with COVID-19, with 11 proteins being increased and 7 proteins decreased in platelets from COVID-19 patients (Figure 2b). Some of the proteins with increased expression in platelets are also enhanced in COVID-19 plasma; G3BP and inter-alpha-trypsin inhibitor heavy chain 3 (ITIH3) and 4 (ITIH4), suggesting the increase may be due to uptake of proteins that are enriched in the plasma. In contrast, α-granule proteins such as VEGFC and thrombospondin-1 (THBS1) were reduced in platelets from COVID-19 patients.

### Reduction in proteins involved in platelet activation or aggregation pathways in COVID-19

37 unique proteins involved in platelet activation or platelet aggregation were detected in the current study. Fifteen of these platelet activation proteins were associated with COVID-19 (Figure 2c). Of these, 12 were reduced in platelets from COVID-19 patients compared to healthy controls, including protein tyrosine kinases (LYN, SYK, JAK2) and Ser/Thr kinases (PKCα, PKCδ and PKCξ). Furthermore, the receptor subunits IX and V, which are part of the platelet GPIb-IX-V complex were also reduced. In contrast, levels of cathepsin G, Apolipoprotein E and interleukin-6 receptor subunit were increased.

### Agonist-induced integrin α_IIb_β_3_ activation is impaired in patients with COVID-19

To relate our proteomic findings to platelet functional responses, we performed flow cytometry studies in PRP to explore activation markers and surface receptors in platelets from COVID-19 patients and controls. Our studies focused on activation of integrin α_IIb_β_3_, the receptor responsible for platelet aggregation, and the α-granule marker P-selectin, which becomes expressed on the surface membrane upon α-granule secretion. Under basal conditions, there was no difference in the activation of integrin α_IIb_β_3_ or surface P-selectin expression between circulating COVID-19 and healthy control platelets (Figure 3a). Platelets from COVID-19 patients showed a small, but significant, reduction in basal levels of the α_IIb_ integrin (CD41) but had similar surface receptor levels for GPVI, GPIbα (CD42b) and β3 integrin (CD61) (Figure 3b). We next performed concentration-response experiments for PAR1-AP, ADP and CRP to evaluate changes in integrin α_IIb_β_3_ activation and P-selectin expression. Our data demonstrates that agonist-induced integrin α_IIb_β_3_ activation is impaired in platelets from COVID-19 patients with a significant reduction of the curve max values, without changes in EC_50_ values (Figure 3c). In contrast to integrin α_IIb_β_3_ activation, agonist-stimulated P-selectin expression was unchanged in platelets from COVID-19 patients (Figure 3d), demonstrating that platelet α-granule secretion remains intact. Similar findings for integrin α_IIb_β_3_ activation and P-selectin expression were observed when using whole blood (Figure 4).

**Figure 3.**
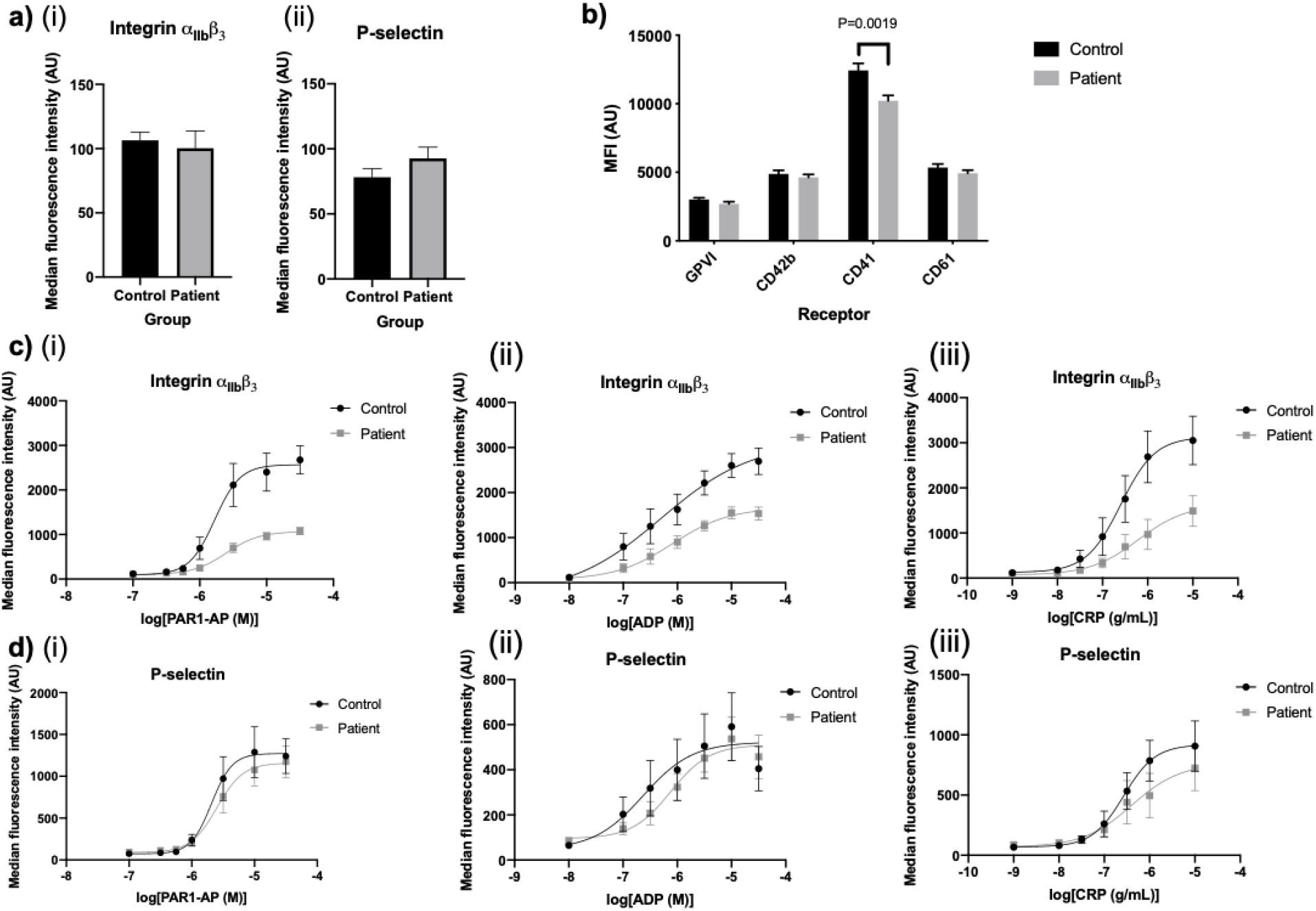
Comparison of integrin α_IIb_β_3_ activation, P-selectin expression and surface receptor levels. a) Comparison of surface receptors and platelet activation markers in diluted platelet rich plasma (PRP) using flow cytometry. Statistical test used is an unpaired t-test unless otherwise indicated (i) Basal integrin α_IIb_β_3_ activation in diluted PRP detected using FITC-PAC1 antibody in COVID-19 patients compared with controls (N=9, Mann-Whitney test, p = 0.18) (ii) Basal P-selectin expression in diluted PRP detected using anti-CD62P-PE antibody in COVID-19 samples compared with controls (N=9, Mann-Whitney test, p = 0.15) b) Basal surface receptor measured in COVID-19 patients and controls: GPVI (N=17-18, p=0.40), CD42b (N=17-18, p=0.47), CD41 (N=17-18, p = 0.0019), CD61 (N=13, p=0.24). c) Concentration-response curves of integrin α_IIb_β_3_ activation with (i) PAR1-AP (Mann-Whitney test comparison of pEC50s, p=0.61, comparison of curve max p = 0.0001), (ii) ADP (comparison of pEC50s, p=0.53, curve max comparison using Mann-Whitney test, p = 0.003) (iii) CRP (comparison of pEC50s, p=0.27, comparison of curve max, p=0.12). d) Concentration-response curves of P-selectin expression in response to increasing concentrations of (i) PAR1-AP (comparison of pEC50s, p=0.76, Mann-Whitney test of comparison of curve max, P=0.96) (ii) ADP (Mann-Whitney test for comparison of pEC50s, p=0.67, Mann-Whitney comparison of curve max, p=0.34) (iii) CRP (comparison of pEC50s, p=0.18, comparison of curve max, p=0.93).

**Figure 4.**
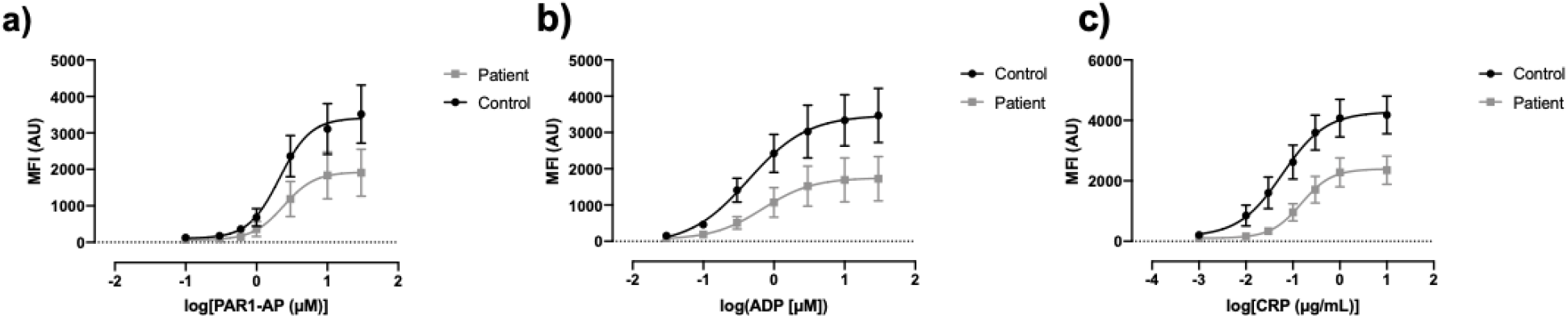
Comparison of integrin α_IIb_β_3_ activation in whole blood. PAC-1 binding using flow cytometry in whole blood (mean ± SEM, N=8) in response to 10-minute stimulation with increasing concentrations of a) PAR1-AP, b) ADP, c) CRP.

### Agonist-induced phosphatidyl serine (PS) exposure is impaired in COVID-19

COVID-19 is associated with hypercoagulopathy and platelets can contribute to coagulation by flipping PS lipids to their outer membranes. We therefore determined platelet procoagulant activity patients by measuring PS exposure in washed platelets using a tagged Annexin V antibody. Under unstimulated conditions, platelets from COVID-19 patients had a small, elevated level of PS exposure compared to controls (2.43 ± 0.32 vs 0.91 ± 0.12, p=0.0001, Figure 5a). In contrast, PS exposure following stimulation with both thrombin and CRP was reduced in the COVID-19 group (35.1 ± 3.1 vs 48.6 ± 4.3, p=0.018, Figure 5b).

**Figure 5.**
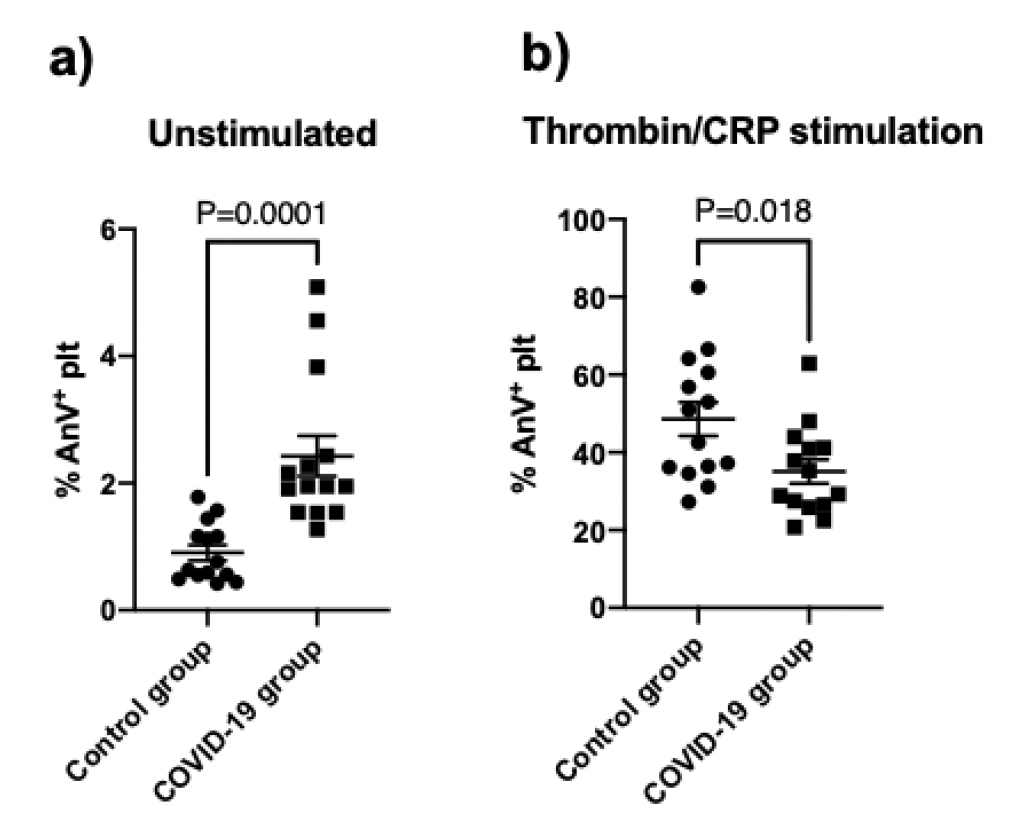
Annexin V binding under basal conditions and after dual stimulation with thrombin and CRP. PS exposure measured by Annexin V binding using flow cytometry in washed platelets at 2 × 10^8^/mL under a) basal conditions (mean ± SEM, N=14) b) stimulation with 1 U/mL thrombin and 10 ug/mL CRP (mean ± SEM, N=14). P values shown for an unpaired student’s t test.

### Platelet-neutrophil interactions are increased in COVID-19

Platelets can contribute to thromboinflammation by interaction with inflammatory cells such as neutrophils. We therefore determined platelet-neutrophil aggregate formation in whole blood from healthy controls and COVID-19 patients by flow cytometry, measuring the platelet marker α_IIb_ (CD41) in the neutrophil gate. Figure 6a demonstrates that 27% (SEM 3%) neutrophils from healthy donors were bound to platelets under basal unstimulated conditions. When control platelets were activated with CRP, 58% (SEM 4%) of the neutrophil population were associated with platelets (Figures 6ai and ii), demonstrating that platelet activation can drive platelet-neutrophil association. In contrast, platelet-neutrophil aggregate formation was already near maximal under unstimulated conditions in the COVID-19 group (72% SEM 5%, p<0.0001 compared to control unstimulated) (Figure 6a). There was no evidence that the proportion of neutrophils bound to platelets further increased (80% (SEM 4%), p>0.05) when stimulated with CRP. To determine the receptors involved in the platelet-neutrophil association, we incubated whole blood for one hour in the presence of a P-selectin CD62P blocking antibody and the α_IIb_β_3_ inhibitor tirofiban. Interestingly, the CD62P blocking antibody significantly reduced basal platelet-neutrophil interactions in blood from COVID-19 patients to the level observed in healthy controls (Figure 6bi). In contrast, the CD62P blocking antibody had no effect on CRP-stimulated platelet-neutrophil interactions in both healthy controls and COVID-19 samples, indicating that additional receptor interactions are likely to take place. Incubation with tirofiban did not prevent or reduce platelet-neutrophil aggregate formation in both healthy controls and COVID-19 patients (Figure 6bi).

**Figure 6.**
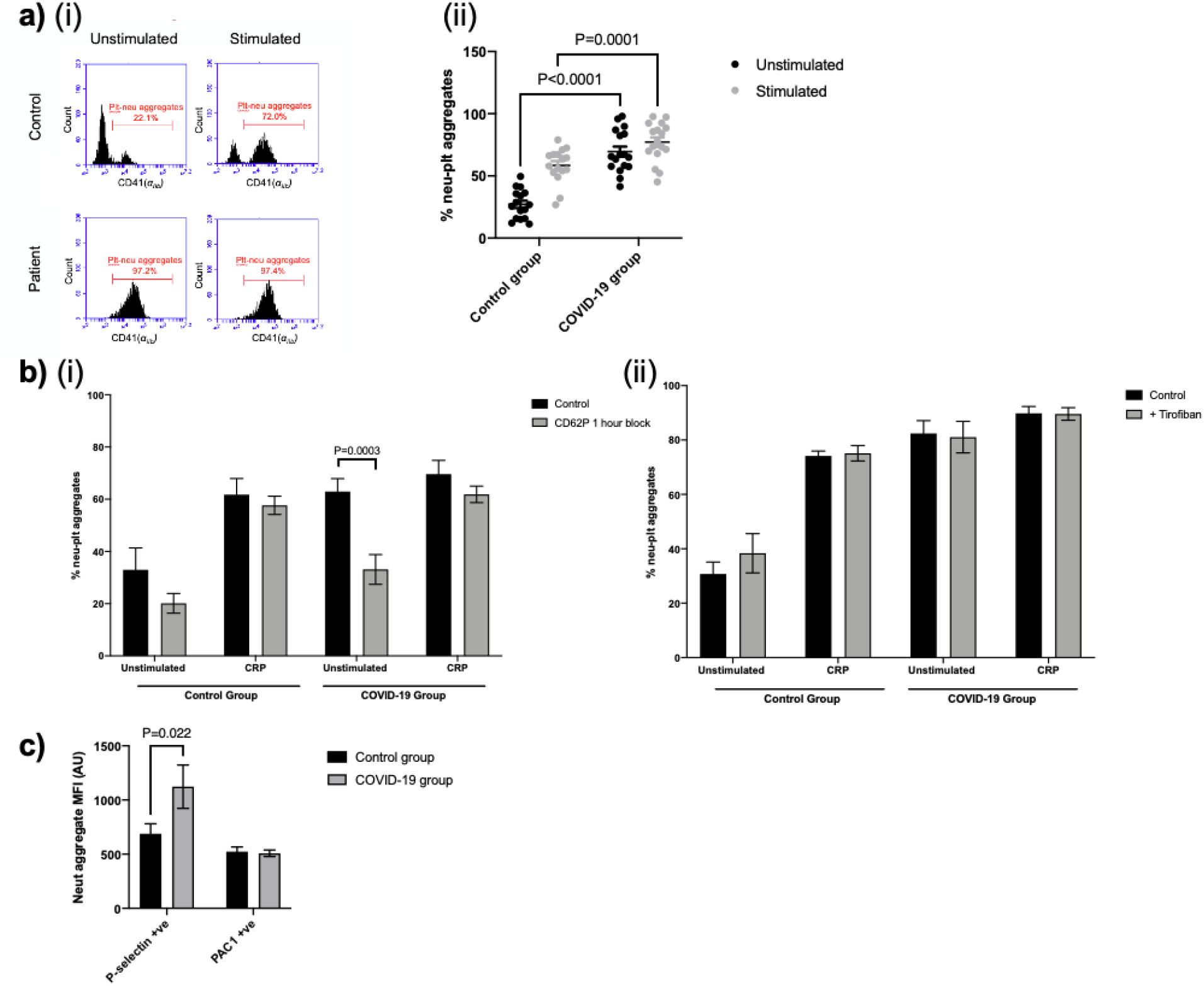
Platelet-neutrophil interactions in patients with COVID-19. a)(i) An example of platelet-neutrophil aggregate formation in whole blood using flow cytometry. Left hand side shows CD41+ events in gated neutrophils under unstimulated conditions and right-hand side following stimulation with 5 μg/mL CRP for 15 mins. (ii) Quantification of the percentage of platelets bound to neutrophils in controls and COVID-19 patients (mean ± SEM, N=17). Control and COVID-19 patient groups compared using an unpaired student’s t-test with P values shown where P<0.05. b) P values indicated in the following figures where P<0.05 (i) Percentage of platelets bound to neutrophils after 1 hour incubation with CD62P blocking antibody (N=4-6, mean ± SEM). Control groups and COVID-19 patient groups compared using a two-way ANOVA with multiple comparisons. b) (ii) Percentage of platelets bound to neutrophils after incubation with tirofiban (mean + SEM, N=7). Results analysed with a mixed effects model with multiple comparisons. c) P-selectin and PAC-1 positive platelets within gated neutrophils (mean ± SEM, N=8). Results analysed with a two-way ANOVA with multiple comparisons.

## Discussion

In this study, we assessed the effect of COVID-19 on the platelet proteome and platelet functional responses. We demonstrate that blood from patients hospitalised with COVID-19 contain at least two platelet populations; free circulating functionally defective platelets and neutrophil-associated platelets. Proteomic studies on the free circulating platelets showed increased levels of a subset of COVID-19 associated plasma proteins and reduced levels of intracellular signalling proteins. These platelets had impaired agonist-mediated responses such as PS exposure and integrin α_IIb_β_3_ activation, whereas P-selectin expression was unaltered. Whole blood analysis indicated that the majority of neutrophils in COVID-19 patients are associated with platelets, an interaction that is dependent on P-selectin. This suggests that platelets drive platelet/neutrophil interaction and immunothrombosis in COVID-19 patients. These main findings are summarised in Figure 7.

**Figure 7.**
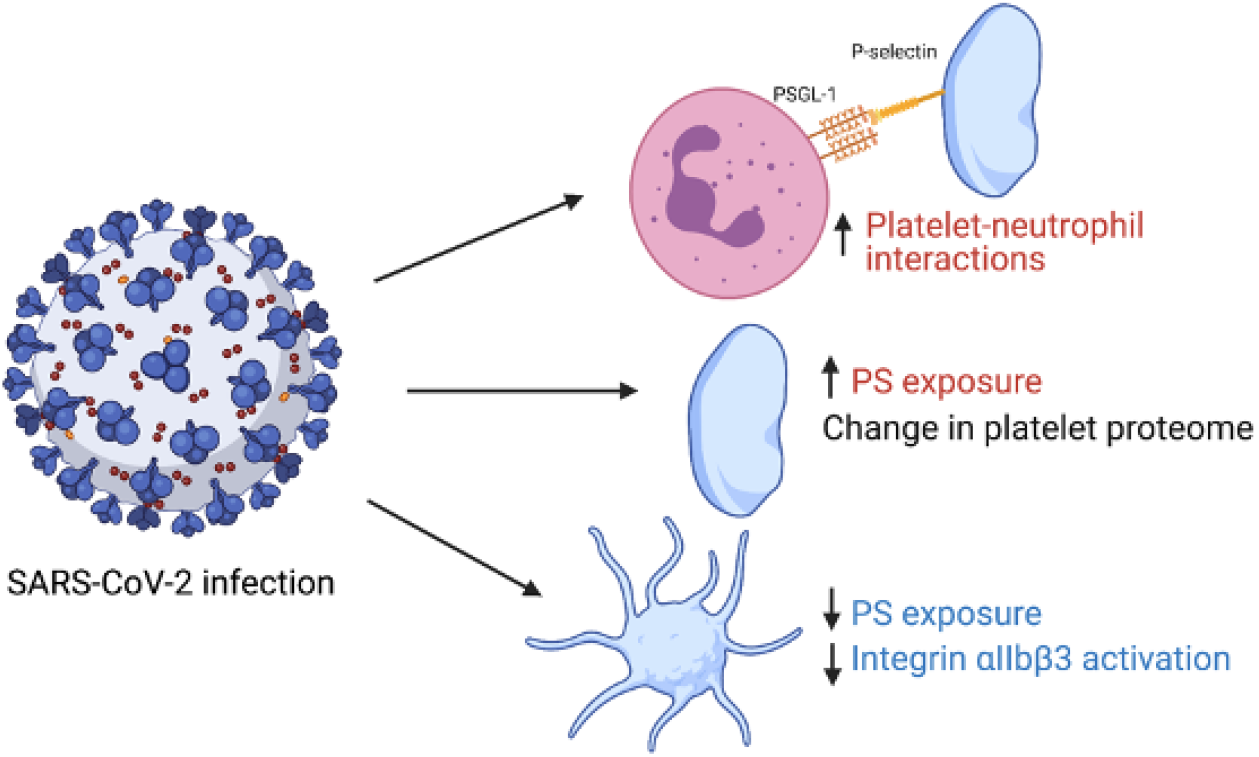
Graphical summary of study findings. Red indicates an increase in variable in patients with COVID- 19 vs healthy controls and blue indicates a decrease. Figure made using biorender.com.

This study showed a large change in the platelet proteome in patients with COVID-19, with strong evidence for altered levels of 858 proteins compared to healthy controls. An interesting finding was the reduction in levels of the TPO receptor c-Mpl in platelets from COVID-19 patients. This reduction is potentially due to enhanced plasma levels of TPO in COVID-19 patients. Increased TPO levels driven, for example by raised IL-6 levels [21, 50], may lead to platelet TPO receptor endocytosis and/or receptor destruction. Conversely, potentially impaired endocytosis of TPO receptors platelets that intrinsically express fewer TPO receptors could also result in a rise in plasma TPO levels. The latter would be consistent with previous platelet transcriptomic findings in COVID-19 patients, reporting a threefold reduction in c-MPL expression [21], which is in the same order of magnitude as the reduction in TPO receptor protein expression levels. We also found reduced protein levels of the signalling molecule JAK2, which constitutively binds to the cytoplasmic tail of the TPO receptor [53]. Both TPO receptor desensitization and internalization, and/or platelets being produced with fewer TPO receptors/JAK2 may thus contribute to the phenotype. Despite the changes in plasma TPO levels and platelet TPO receptor/JAK2 levels, we and others found that platelet counts are in the normal range in COVID-patients [21, 54]. Thus, the alterations do not lead to significant alterations in platelet count, although we cannot rule out that any potential changes are masked by platelet association with immune cells or dexamethasone treatment of COVID-19 patients [55].

We also compared our platelet proteomic findings to three recently reported studies on changes in the plasma/serum proteome in COVID-19 patients [42–44]. Using a threshold setting of the protein being detected in two out of the three published studies in plasma/serum, we found a subset of the proteins that were highly enriched in platelets from COVID-19 patients. Proteins that were more than fourfold enriched in platelets include the acute phase proteins; C-reactive protein, SAA1, LBP and G3BP. We also found an enrichment of proteins related to the complement system (complement component C9 and C4b-binding protein alpha chain (C4BPA)), Inter-alpha-trypsin inhibitor heavy chain H (ITIH3 and ITIH4), and the leukocyte marker CD14. Interestingly, two proteins that were downregulated in COVID-19 plasma, were upregulated in platelets: Apolipoprotein A-I (ApoA1) and Inter-alpha-trypsin inhibitor heavy chain H1. ApoA1 plasma levels are inversely associated with COVID-19 severity, suggesting a protective function [56, 57]. For the plasma/serum proteins which were associated with COVID-19 and also associated with COVID-19 in platelets, a transcriptomic study did not find a difference in levels of transcripts for these genes [21], suggesting platelet protein uptake from plasma, rather than an alteration in megakaryocyte gene expression is responsible for enhanced levels. One exception is G3BP, whose transcript is upregulated in platelets from COVID-19 patients [21]. Platelet-derived G3BP may therefore contribute to the observed increase in plasma G3BP levels in COVID-19 patients. Of note, G3BP and its receptor/ligand galectin-3 have been reported to contribute to platelet hyperactivity and venous thrombosis [58, 59].

When assessing proteins involved in platelet secretion and/or granule content, we also found proteins that were COVID-19 plasma biomarkers (G3BP, ITIH3 and ITIH4) and acute phase proteins. Interestingly, proteins known to be released upon α-granule secretion, such as thrombospondin-1 and VEGF-C were reduced in COVID-19 platelets, suggesting granule markers are reduced, potentially pointing to platelet pre-activation/exhaustion. The latter is also reflected by a reduction in a range of signalling proteins detected in platelets from COVID-19 platelets. However, we did not find evidence of platelet pre-activation, as basal integrin α_IIb_β_3_ and P-selectin expression in platelets from COVID-19 patients were unaltered. There was a small increase in PS exposure, although this study may have been underpowered to detect a subtle increase. Although some previous studies have suggested platelet hyperactivity in COVID-19 patients [25, 32, 60, 61], we observed that integrin α_IIb_β_3_ activation in response to platelet agonists was impaired, whereas P-selectin expression was unaltered, confirming previous observations in response to collagen [21, 25]. Similarly, we found that agonist stimulated PS exposure was attenuated in the COVID-19 group, in agreement with previous findings [62]. Both the proteomics and pathway analysis pointed towards impaired platelet signaling, with a reduction in levels of PKCα, and inhibition of Tec kinase and phospholipase C. Impairment in these signaling pathways may underlie the observed reduction in platelet function. Furthermore, a small but significant reduction in levels of CD41 (α_IIb_) expression in COVID-19 patients may further contribute to the measured impairment in agonist-induced integrin α_IIb_β_3_ activation. Interestingly, as previous studies found normal or enhanced platelet aggregation in COVID-19 patients [21, 63], the residual integrin α_IIb_β_3_ activation may still be sufficient for normal aggregation in the COVID-19 group.

The plasma protein S100A8/A9 did not pass our selection criteria of being elevated in two out of three proteomic studies [42–44], however several studies have reported raised plasma S100A8/A9 in COVID-19 patients and plasma levels correlated to poor outcome [64–67]. S100A8/A9 can induce platelet procoagulant responses and promote platelet/neutrophil interaction by enhancing P-selectin levels [66]. Here, we found that S100A8/A9 protein levels were enhanced by more than 4-fold (log_2_ fold change of 2.25) in platelets in COVID-19 patients. As platelet transcript levels in COVID-19 patients were unchanged [21], this is likely to be the result of enhanced platelet binding and uptake of plasma S100A8/A9, potentially through GPIb/CD36 receptor interaction [66]. The enhanced basal platelet PS exposure in COVID-19 patients detected in our study would be consistent with this.

In whole unstimulated blood, patients with COVID-19 exhibited platelet-neutrophil interactions, which is consistent with previous studies reporting enhanced platelet-leukocyte interactions in COVID-19 [21, 25, 32]. The majority of neutrophils in COVID-19 patients were associated with platelets. Platelet-neutrophil interactions were found to be dependent on P-selectin and not integrin α_IIb_β_3_. This agrees with our finding that neutrophil-associated platelets were positive for P-selectin but lacked activated integrin α_IIb_β_3_. The latter may be explained by the reversibility of integrin α_IIb_β_3_ activation, where initial platelet activation leads to both P-selectin expression and integrin α_IIb_β_3_ activation, but integrin α_IIb_β_3_ then reverts to a low affinity state [68, 69]. However, we cannot rule out that COVID-19 conditions may induce platelet P-selectin expression in the absence of integrin α_IIb_β_3_ activation, such as has recently observed in a small fraction of CRP/PAR-1 activated platelets [70]. The recent study by Colicchia et al [66] is also of interest as they demonstrated that S100A8/A9 can induce P-selectin expression whilst inducing an integrin α_IIb_β_3_ activatory state that lacked aggregatory ability and had low-fibrinogen binding capacity.

One mechanism by which COVID-19 may affect platelet function is by direct modulation by SARS-CoV-2 [71]. Most studies were unable to detect platelet expression of the SARS-CoV-2 entry receptor ACE2 at both mRNA and protein level [21, 23], although a few studies reported ACE-2-mediated regulation of platelet function [36]. Data from our proteomics study support the lack of ACE2 expression in human platelets. We cannot exclude that there are alternative direct modes of regulation, such as through CD147 (basigin) [48], which we here confirm is expressed on platelets. We did however not find evidence for viral protein expression in platelets from COVID-19 patients, suggesting that SARS-CoV-2 cannot enter and/or replicate in human platelets.

There are a few limitations which are important to note. As the vast majority of patients were on dexamethasone and heparin, we cannot rule out that these medications may contribute to the platelet phenotype we have observed. However, as this was the recommended care for those hospitalized with COVID-19, it was challenging to specifically recruit participants who were not on these medications. Secondly, the healthy controls were younger than the recruited COVID-19 patients, and on average had a lower body mass index (BMI). Therefore, we cannot rule out confounding from variables such as BMI. Finally, the control participants self-reported being negative for SARS-CoV-2. It is possible that control participants could have been carrying the virus and were asymptomatic. However, we would expect that this would only result in smaller differences in platelet function between groups.

Overall, our data suggests the presence of two platelet populations in patients with COVID-19. The first is circulating platelets with an altered proteome, increased basal PS exposure and reduced agonist-induced integrin α_IIb_β_3_ activation. The second platelet population is P-selectin expressing neutrophil-associated platelets. Furthermore, circulating platelets from COVID-19 patients have a unique protein signature, with multiple COVID-19 associated plasma proteins being markedly enhanced. Our data shows a complex picture and suggests that platelet driven thromboinflammation may be one of the key drivers enhancing the risk of thrombosis in COVID-19 patients. The data also point towards potential mechanisms of this effect, which now need to be further characterised.

## Acknowledgements

This work was supported by the University of Bristol Elizabeth Blackwell Institute, the University of Bristol Alumni, the Southmead Hospital Charity, the Welcome Trust (216277/Z/19/Z, 219472/Z/19/Z) and the British Heart Foundation (FS/17/60/33474, RG/15/16/31758, PG/17/62/33190, PG/21/10760, SP/F/21/150023). LJG is supported by the BHF accelerator award AA/18/1/34219.

We thank Borko Amulic and Christopher Rice for their helpful discussions over the observed interactions between platelets and neutrophils. This research was funded in whole, or in part, by the Wellcome Trust. For the purpose of Open Access, the author has applied a CC BY public copyright licence to any Author Accepted Manuscript version arising from this submission.

## Authorship Contributions

IH obtained ethical approval and led the study. LJG, CMW and IH designed experiments. FH, DA and AM were involved in obtaining patient samples and patient information. LJG, CMW, JK and KLB performed experiments. PAL and KJH generated the proteomics results. LJG, KLB and IH analysed and interpreted the proteomic data. AD supported interpretation of proteomic results. LJG and IH wrote the manuscript. AWP and SJM contributed to discussion. All authors reviewed and/or edited the manuscript.

## Conflicts of Interest

All authors declare no conflict of interest.

## Notes

### Competing Interest Statement

The authors have declared no competing interest.

