## Supplementary Methods for "Alterations in platelet proteome signature and impaired platelet integrin α_IIb_β_3_ activation in patients with COVID-19"

*Materials*

Protease-activated receptor 1 (PAR-1)-activating peptide (SFLLRN-NH2) was from Bachem (Bubendorf, Switzerland), crosslinked collagen-related peptide (CRP-XL) was from Peptide Synthetics. Adenosine diphosphate (ADP) was from Sigma-Aldrich (Poole, UK). cOmplete mini protease inhibitor tablets and phosSTOP phosphatase inhibitors were from Roche Life Sciences (Welwyn Garden City, UK). The Pierce bicinchoninic acid (BCA) assay and the P-selectin (CD62P) blocking antibody was from ThermoFisher Scientific (Altrincham, UK). Sodium Citrate Vacutainer® tubes, FixLyse, FITC-conjugated Annexin V, PE-Cy5-conjugated anti-CD42b, FITC-conjugated CD61 antibody, Fc block, PE-conjugated human platelet GPVI antibody, PAC1-FITC, anti-CD62P-PE, PKA RIIβ and TPO-receptor antibodies were from BD (Wokingham, UK). FITC-conjugated anti-human CD41 and CD42b antibody were from BioLegend (London, UK). PKCα antibody was obtained from Cell Signalling Technology (New England Biolabs, Hitchin, UK) and was from Invitrogen

*TMT protein quantification*

TMT Labelling and High pH reversed-phase chromatography: Aliquots of 50µg of each sample were digested with trypsin (2.5µg trypsin per 100µg protein; 37°C, overnight), labelled with Tandem Mass Tag (TMT) eleven plex reagents according to the manufacturer’s protocol (Thermo Fisher Scientific, Loughborough, UK) and the labelled samples pooled. A 50ug aliquot of the pooled sample was evaporated to dryness, resuspended in 5% formic acid and then desalted using a SepPak cartridge according to the manufacturer’s instructions (Waters, Milford, Massachusetts, USA). Eluate from the SepPak cartridge was again evaporated to dryness and resuspended in buffer A (20 mM ammonium hydroxide, pH 10) prior to fractionation by high pH reversed-phase chromatography using an Ultimate 3000 liquid chromatography system (Thermo Scientific). In brief, the sample was loaded onto an XBridge BEH C18 Column (130Å, 3.5 µm, 2.1 mm X 150 mm, Waters, UK) in buffer A and peptides eluted with an increasing gradient of buffer B (20 mM Ammonium Hydroxide in acetonitrile, pH 10) from 0-95% over 60 minutes. The resulting fractions (15 in total) were evaporated to dryness and resuspended in 1% formic acid prior to analysis by nano-LC MSMS using an Orbitrap Fusion Lumos mass spectrometer (Thermo Scientific).

Nano-LC Mass Spectrometry: High pH RP fractions were further fractionated using an Ultimate 3000 nano-LC system in line with an Orbitrap Fusion Lumos mass spectrometer (Thermo Scientific). In brief, peptides in 1% (vol/vol) formic acid were injected onto an Acclaim PepMap C18 nano-trap column (Thermo Scientific). After washing with 0.5% (vol/vol) acetonitrile 0.1% (vol/vol) formic acid peptides were resolved on a 250 mm × 75 μm Acclaim PepMap C18 reverse phase analytical column (Thermo Scientific) over a 150 min organic gradient, using 7 gradient segments (1-6% solvent B over 1min., 6-15% B over 58min., 15-32%B over 58min., 32-40%B over 5min., 40-90%B over 1min., held at 90%B for 6min and then reduced to 1%B over 1min.) with a flow rate of 300 nl min−1. Solvent A was 0.1% formic acid and Solvent B was aqueous 80% acetonitrile in 0.1% formic acid. Peptides were ionized by nano-electrospray ionization at 2.0kV using a stainless steel emitter with an internal diameter of 30 μm (Thermo Scientific) and a capillary temperature of 300°C.

All spectra were acquired using an Orbitrap Fusion Lumos mass spectrometer controlled by Xcalibur 3.0 software (Thermo Scientific) and operated in data-dependent acquisition mode using an SPS-MS3 workflow. FTMS1 spectra were collected at a resolution of 120 000, with an automatic gain control (AGC) target of 200 000 and a max injection time of 50ms. Precursors were filtered with an intensity threshold of 5000, according to charge state (to include charge states 2-7) and with monoisotopic peak determination set to peptide. Previously interrogated precursors were excluded using a dynamic window (60s +/-10ppm). The MS2 precursors were isolated with a quadrupole isolation window of 0.7 m/z. ITMS2 spectra were collected with an AGC target of 10 000, max injection time of 70ms and CID collision energy of 35%.

For FTMS3 analysis, the Orbitrap was operated at 50 000 resolution with an AGC target of 50 000 and a max injection time of 105ms. Precursors were fragmented by high energy collision dissociation (HCD) at a normalised collision energy of 60% to ensure maximal TMT reporter ion yield. Synchronous Precursor Selection (SPS) was enabled to include up to 10 MS2 fragment ions in the FTMS3 scan.

*Data analysis:* The raw data files were processed and quantified using Proteome Discoverer software v2.1 (Thermo Scientific) and searched against the UniProt Human database (downloaded August 2020; 167789 sequences) using the SEQUEST HT algorithm. Peptide precursor mass tolerance was set at 10ppm, and MS/MS tolerance was set at 0.6Da. Search criteria included oxidation of methionine (+15.995Da), acetylation of the protein N-terminus (+42.011Da) and Methionine loss plus acetylation of the protein N-terminus (-89.03Da) as variable modifications and carbamidomethylation of cysteine (+57.021Da) and the addition of the TMT mass tag (+229.163Da) to peptide N-termini and lysine as fixed modifications. Searches were performed with full tryptic digestion and a maximum of 2 missed cleavages were allowed. The reverse database search option was enabled, and all data was filtered to satisfy false discovery rate (FDR) of 5%.
